# Context-dependent differences in the functional responses of Lactobacillaceae strains to fermentable sugars

**DOI:** 10.1101/2022.07.11.499608

**Authors:** Ronit Suissa, Rela Oved, Harsh Maan, Omri Gilhar, Michael Meijler, Omry Koren, Ilana Kolodkin-Gal

## Abstract

Lactobacillaceae are Gram-positive rods, facultative anaerobes, and belong to the lactic acid bacteria (LAB) that frequently serve as probiotics. We systematically compared five LAB strains for the effects of different carbohydrates on their free-living and biofilm lifestyles. We found that fermentable sugars triggered a heterogeneous response in LAB strains, frequently manifested specifically in altered carrying capacity during planktonic growth and colony development. The fermentation capacities of the strains were compatible and could not account for heterogeneity in their differential carrying capacity in liquid and on a solid medium. Among tested LAB strains, *L. paracasei, and L. rhamanosus GG* survived self-imposed acid stress while *L. acidophilus* was extremely sensitive to its own glucose utilization acidic products. The addition of a buffering system during growth on a solid medium significantly improved the survival of most tested probiotic strains during fermentation. We suggest that the optimal performance of the beneficial microbiota members belonging to lactobacilli is heterogeneous and varies as a function of the growth model and the dependency on a buffering system.

## Introduction

Firmicutes are dominant phylum in the human microbiota and the proportions of these phylum members vary between individuals and are greatly influenced by the local pH(1). Accordingly, the stomach is the least diverse growth niche within the human gastrointestinal tract (due to its extreme acidity). In the microbiome, the presence of probiotic strains varies between healthy individuals. Probiotics are defined simply as “microorganisms that when administered in adequate amounts confer a health benefit to the host” (2). Probiotic strains are consumed either as fresh fermentation products or as dried bacterial supplements with Lactobacillaceae and Bifidobacteria, being the two widely used probiotic species(3, 4). The main strains currently used for probiotic formulation were originally isolated from fermented products or humans(5–7).

Lactobacillaceae are Gram-positive rods, facultative anaerobes, and belong to the lactic acid bacteria (LAB) group, as lactic acid is their main end-product of carbohydrate metabolism(8, 9). This family is naturally found in the gastrointestinal tract (GIT) of humans and animals as well as in the urogenital tract of females(10). LAB are considered efficient fermenters, proficient in the production of energy under anaerobic conditions, or when oxygen is limited. The fermentation process involves the oxidation of carbohydrates to generate a range of products including organic acids, alcohol, and carbon dioxide(11). In general, the response of all LAB strains to fermentation is considered to be uniform and primarily depends the capacity to utilize glucose and its products (2).

In LAB, a phosphotransferase system (PTS) transports glucose across the membrane that is then used by the glycolysis pathway to produce pyruvate. This pathway generates energy and consumes NAD^+^. Pyruvate is then converted into L- and D-lactate by the stereospecific NAD-dependent lactate dehydrogenases (LDHs), LdhL and LdhD, respectively, which regenerate NAD^+^ and maintain the redox balance(12). In the presence of oxygen and following the depletion of glucose, lactate is oxidized to pyruvate via the Lactate Oxidase enzyme followed by the production of acetate using pyruvate oxidase and ACK enzymes. The formation of acetate as the major fermentation end product, results in homoacetic fermentation (13). In addition to the importance of glucose utilization to microbial metabolism, glucose and fructose induce biofilm formation in *lactobacillus rhamnusos GG*, the most characterized LGG strain due to changes in the protein abundance and major changes in the surface proteome, indicating that changes in carbon sources and their concentrations induce more complex adaptations than alterations in microbial growth (14). Whether these carbohydrate utilization induces regulatory effects in additional LAB strains remain to be determined.

To map adaptations to carbohydrates that are growth dependent and independent, we systematically compared the response to metabolic stress in microaerophilic (CO_2_ enriched environments) >3% CO^2^) conditions of the five probiotic Lactobacillaceae species: *Lacticaseibacillus rhamnosus GG* (15), *Lacticaseibacillus casei* (16), *Lactobacillus acidophilus* (17), *Lacticaseibacillus paracasei*, and *Lactiplantibacillus plantarum* (18) during planktonic growth and colony biofilm formation. *Bacillus coagulans*, a probiotic *Bacilli* belonging to the same phylum(19), was studied as a non-Lactobacillaceae LAB control strain. Under our conditions, similar glucose utilization efficiencies were observed between the species (as judged by the levels of the end products: lactate and acetate). Our results indicate that a differential response to carbohydrates is correlated with a differential acid tolerance, rather than differential production of organic acids. Thereof, LAB bacteria significantly differ in their response to their own self-imposed acid stress from glucose utilization products and can be clustered into fermentation resistant and fermentation sensitive strains. The differential adaptation to acidic products included reversible changes in cellular organization and colony formation.

## Results

### Fermentable sugars specifically but heterogeneously affect the carrying capacity during planktonic growth

To test whether fermentable sugars affect growth differently than non-fermentable sugars, we grew five probiotic Lactobacillaceae species on a rich medium (TSB) in a shaking culture either with glucose (fermentable) or with raffinose (non-fermentable). All bacteria grew similarly in rich medium. The addition of glucose induced growth as judged by an increased carrying capacity of *L. acidophilus* and *LGG* compared to TSB alone, but its effect on growth *in L. paracasei, L. casei L. plantarum and B*.*coagulans* was extremely noisy (Figure 1). To better distinguish between growth patterns of the different probiotic strains, we further analyzed the growth with the software GrowthRates 3.0. We extracted the length of the lag phase, the growth rate and the carrying capacity of all LAB strains with the different treatments (20). Our results indicated that the heterogeneous response is primarily manifested in an altered carrying capacity (e.g.: the maximal OD of the cultures) (Figure 2A). Strains showed varied carrying capacity from each other, and indeed *L. acidophilus* and *LGG* had enhanced carrying capacity in response to Glucose, while other probiotic strains did not exhibit significant response. In all species the growth rate and lag time were not significantly altered by glucose addition (Figures 2B and C).

**Figure 1.**
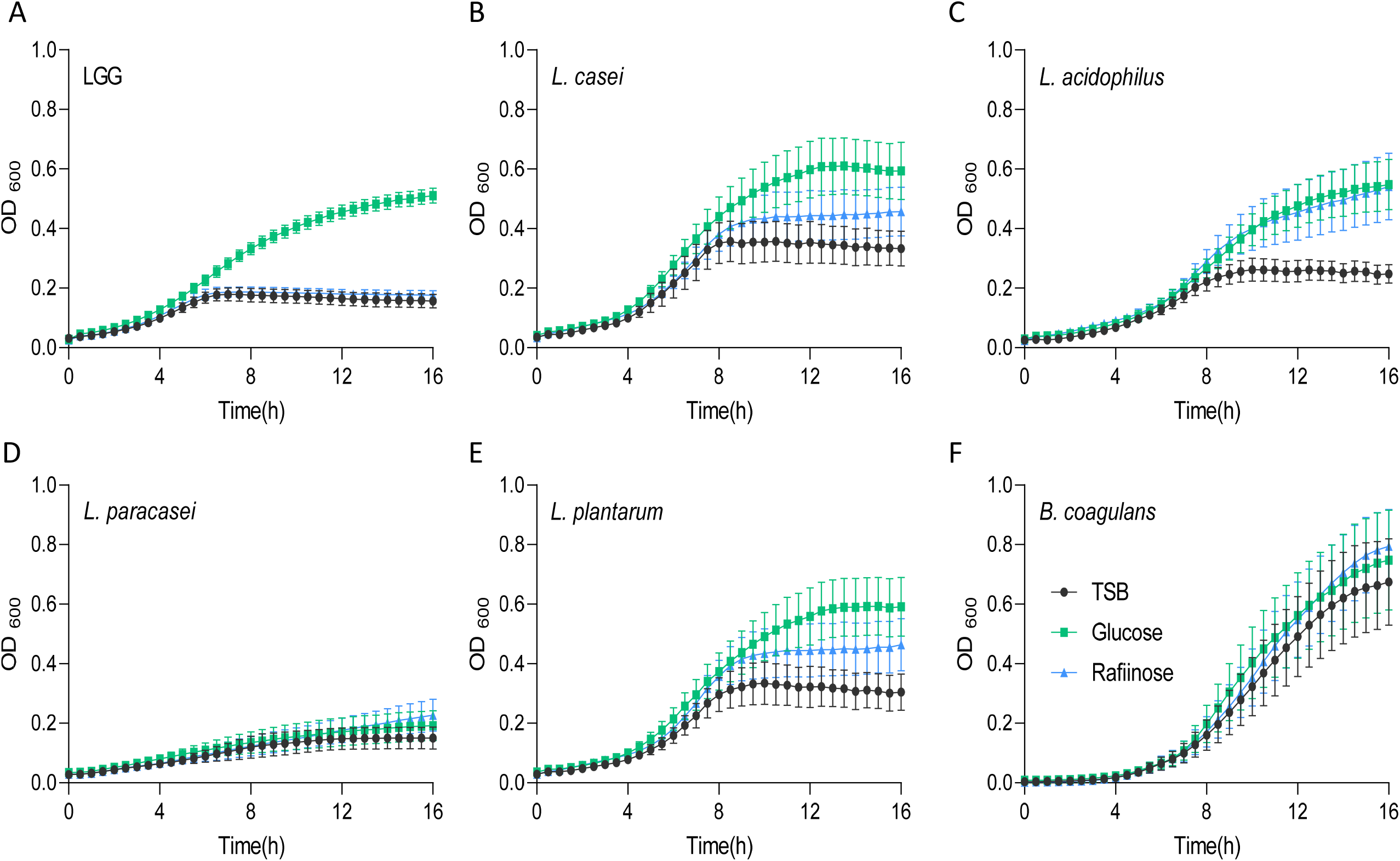
Heterogeneous alterations in planktonic growth of LAB strains during glucose utlization. Planktonic growth of the indicated species **(A)** LGG **(B)** *L. casei* **(C)** *L. acidophilus* **(D)** *L. paracasei* **(E)** *L. plantarum* **(F)** and *B. coagulans* in TSB medium (control) in TSB medium either supplemented or not with glucose (1% W/V) and raffinose (1% W/V). Graphs represent mean ± SEM from six independent experiments (n = 24).

**Figure 2.**
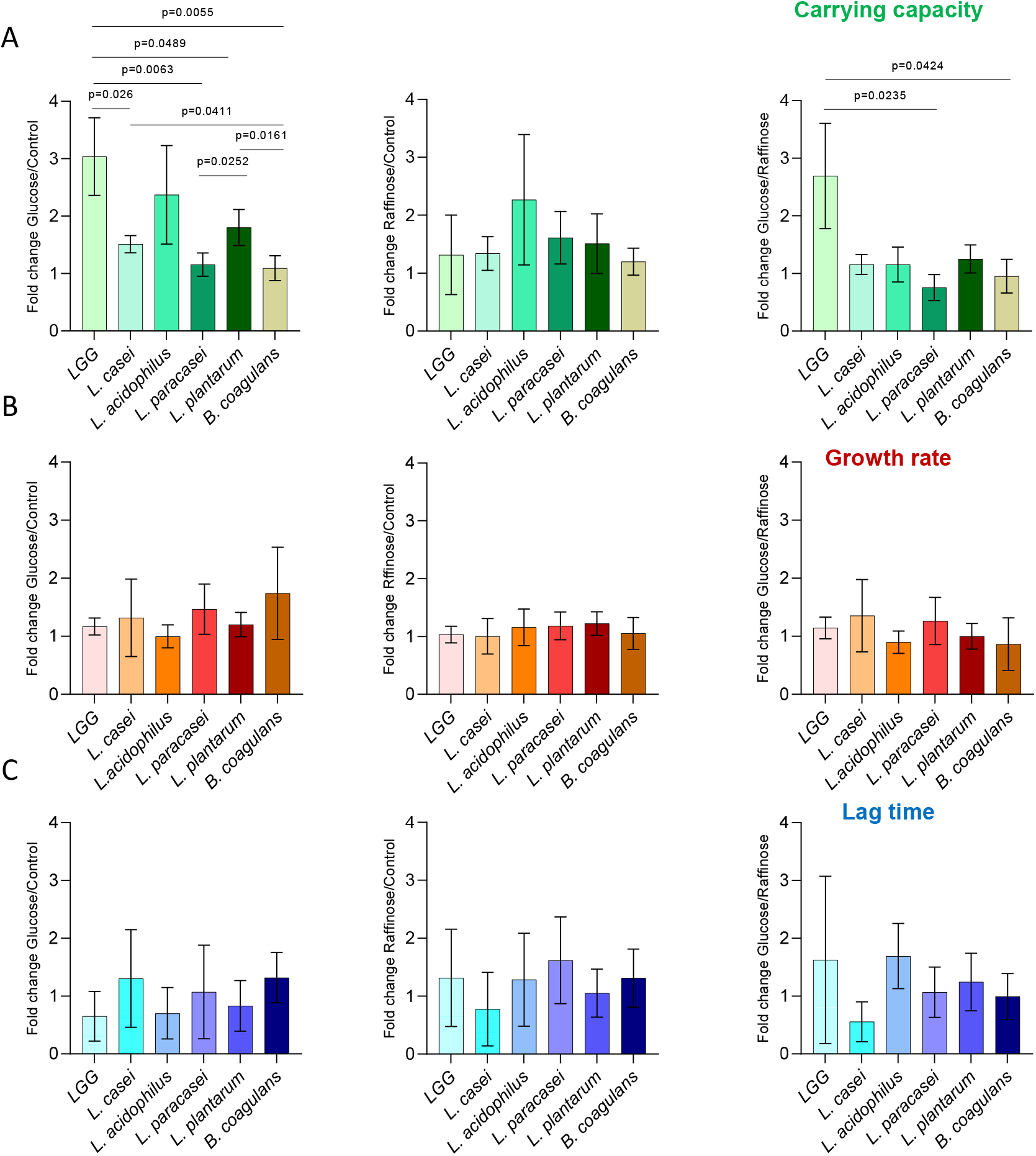
Carbon-dependent growth induction is specifically manifested by an altered carrying capacity. Analysis of planktonic growth from the data shown in Figure 1 using GrowthRates 3.0. **A)** Fold Change in carrying capacity of indicated species between glucose/control, raffinose/control and glucose/raffinose. **B)** Fold Change in growth rate of indicated species between glucose/control, raffinose/control and glucose/raffinose. **C)** Fold Change in lag time of indicated species between glucose/control, raffinose/control and glucose/raffinose. Graphs represent mean ± SD from six independent experiments (n = 24). Statistical analysis was performed using Brown-Forsthye and Welch’s ANOVA with Dunnett’s T3 multiple comparisons test. P < 0.05 was considered statistically significant.

We then asked whether the induction of growth is an outcome of fermentable sugar utilization, as fermentation is a metabolic process allowing the production of energy under anaerobic conditions. To answer that question, we assessed the growth of the bacteria on a medium supplemented with raffinose, a sugar source less compatible with fermentation, as it was indicated that alpha-galactosidase, responsible for the hydrolysis of this sugar, is not enzymatically active (21–23). Raffinose did not induce the growth of LGG, suggesting that the induction of growth depends on the fermentable nature of the sugar (Figures 1 and 2A). In contrast, the carrying capacity of *L. acidophilus* cultures increased with both glucose and raffinose (Figure 1) but failed to meet statistical analysis of each parameter of growth (Figures 2B and C). In general, the addition of raffinose failed to induce significant alterations in growth parameters in all strains. Similar results were observed when the bacteria were grown with mannose (fermentable) that induced an enhanced carrying capacity and xylose non-fermentable), failing to induce significant alterations of growth (Figure S1). With the exception of carrying capacity, other parameters of growth remained unaltered by sugar application In all strains (Figures 1, 2 and S1).

### Differential response of Lactobacillaceae to fermentation is not a readout of organic acid production

To compare the fermentation efficiency, we measured the acidity of the medium before and after growth in the presence and absence of glucose. Indeed, after 24h the pH of the medium decreased in both glucose and raffinose (Figure 3A). The acidity of Lactobacillaceae conditioned medium grown in the absence of glucose was approximately 5, while the addition of glucose to the medium lowered the pH to 4 or less confirming that upon application of glucose, the acidification of the growth media was enhanced in all Lactobacillaceae that were tested. Compared with other tested strains, *B. coagulans* altered the pH more modestly, e.g.: to 4.5 in the presence of glucose and 6 in its absence (Figure 3A). The pH drop of the growth medium source was comparable between all Lactobacillaceae stains in glucose and raffinose, except in LGG which was inert to raffinose. This result indicates that differential utilization of glucose may not account for the differential carrying capacities of the cultures in the presence of glucose.

**Figure 3.**
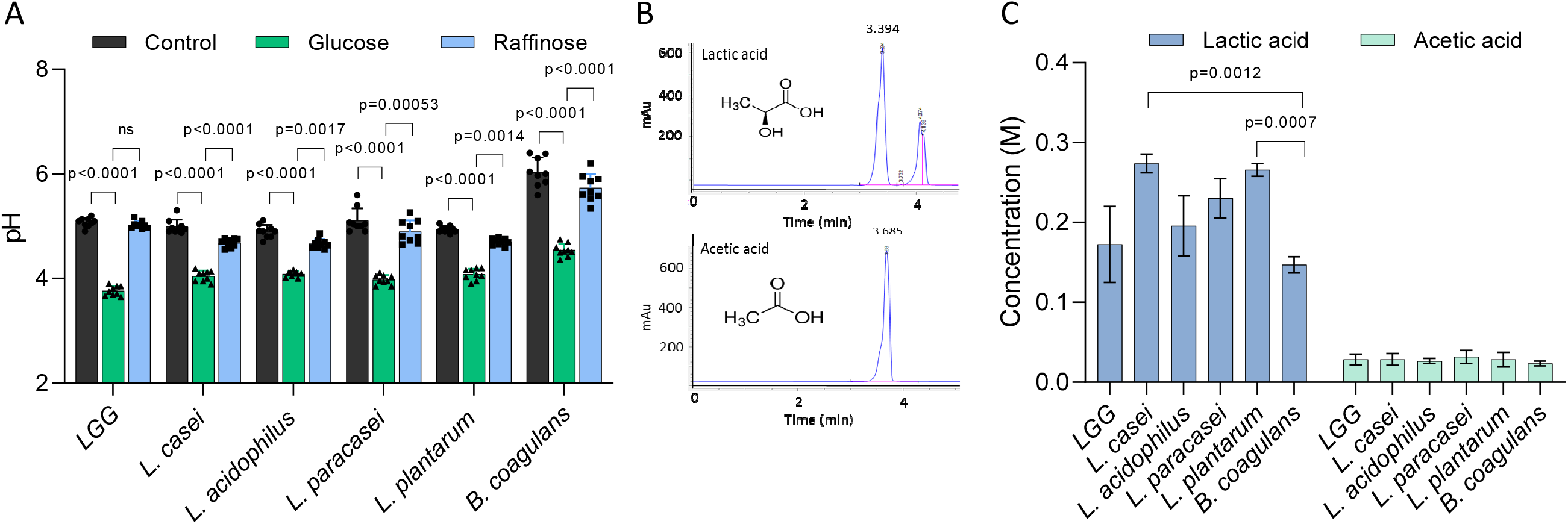
The variation at the specie level in growth enhancement is not due to glucose catabolism. **A)** The pH of the conditioned media of indicated species in TSB medium (control) and TSB medium supplemented with glucose (1% W/V) and raffinose (1% W/V). Statistical analysis was performed using two-way ANOVA followed by Dunnett’s multiple comparison test. P < 0.05 was considered statistically significant. **B)** HPLC chromatograms indicates an exclusive peak and stable retention time for the pure standards of Lactic acid and Acetic acid. **C)** The concertation of lactic acid and acetic acid produced by the indicated species. Bacteria were grown on MSgg medium supplemented with glucose (1% W/V) for 24h. The conditioned medium from the cultures was collected and analyzed using C-18 column. Statistical analysis was performed using Brown-Forsthye and Welch’s ANOVA with Dunnett’s T3 multiple comparisons test. P < 0.05 was considered statistically significant.

In order to confirm that the uniform drop in pH levels results from the differential accumulation of organic acids among the tested LAB species, we measured the accumulation of glucose utilization products (organic acids), focusing on the levels of key organic acids, lactate and acetate using High-performance liquid chromatography (HPLC) (calibrated as in Figure S2). In general, lactic acid was ten times more abundant than acetic acid in the growth media. When we statistically analyzed the concentration of lactate produced by the bacteria, there was no single species that was significantly different from the other group members, indicating that the production of organic acids and thereby glucose utilization is similarly performed under our conditions (Figure 3B and C) and cannot account for the differential carrying capacity.

To exclude that the differential response to fermentation is a result of different enzymatic activities of enzymes involved in the process, we studied *in silico* the presence of lactate dehydrogenase from different LAB strains. Alignment of LDH proteins from different Lactobacillaceae (Figure S3) indicated that *LGG L. paracasei* and *L. casei* have two groups of homologues LDH proteins with more than 90% identity. 16S alignment showed that all species have at least 89% similarity, and the evolutionary closest species are *LGG L. casei* and *L. paracasei* based on 16S. Therefore, the key differences revealed by growth analysis on fermentable sugars are despite the high evolutionary closeness of all five species.

### Self-imposed acid stress in biofilm colonies of LAB species

To further study the differential response to fermentation in Lactobacillaceae, we examined the growth on solid medium under static conditions, which is frequently correlated with biofilm formation(24). To test whether the addition of fermentable sugars plays a role when grown on solid media under microaerophilic conditions, we assessed the colony morphology with the addition of glucose and raffinose (Figure 4A). In all strains, the application of glucose-induced the formation of asymmetric, smaller, and morphologically different colonies. Raffinose did not induce the same morphological changes in the colony structure, suggesting glucose metabolism has a role also in shaping the colony architecture (Figure 4A). Solid growth media from bacteria grown on TSB or TSB with raffinose had similar pH values of approximately 5.5 (Figure 4B). The addition of glucose to the growth media lowered the pH significantly suggesting that glucose-induced fermentation in all tested strains under these conditions. To test whether acid stress is related to the alterations in colony morphology we monitored cell death in the presence and absence of a buffering system. As shown, in all strains, with the exceptions of *L. paracasei*, the buffering system significantly enhanced the survival of the cells, and acid-dependent cell death during fermentation significantly varied between the strains (Figure 4C-4D). The addition of a buffer also resulted in colony morphology comparable to the morphology of strains grown without glucose. While cell death was reduced, a buffer did not fully restore the viability of the cells within a colony, it was sufficient to restore colony morphologies to those observed in a glucose-free solid medium (Figure 4A). These results may suggest that cell death on solid biofilm media primarily but not solely results from differential acid sensitivity, and that adaptation to acidic stress triggers alterations in biofilm formation capacities independently of cell counts. These results, observed in all LAB strains are consistent with recent findings that the stress protein Hsp plays an important role in shaping colony morphology under acidic pH in a single specie: *L. plantarum* (25).

**Figure 4.**
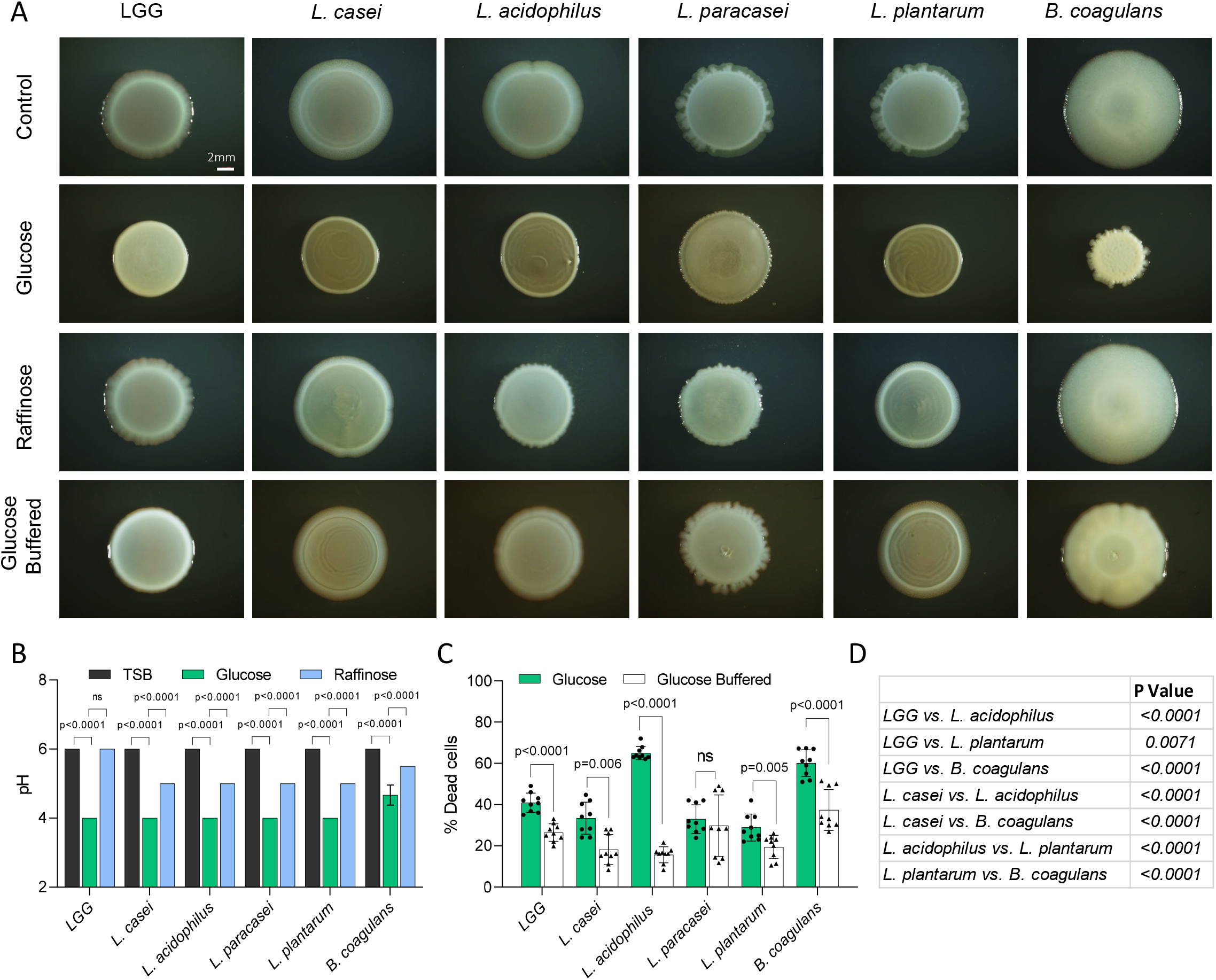
Self-imposed acidic stress has a role in shaping microbial colonies. **A)** Shown are the indicated biofilm colonies grown on TSB medium (control), TSB medium supplemented with glucose and raffinose (1% W/V), and TSB medium supplemented with glucose (1% W/V) + buffer. Biofilms were grown at 37° C in in CO_2_ enriched environment. Biofilms colonies were imaged at 72 hours post inoculation. Scale bar = 2mm. **B)** Measurement of pH of the indicated species as shown in A. Statistical analysis was performed using two-way ANOVA followed by Dunnett’s multiple comparison test. P < 0.05 was considered statistically significant. **C)** Flow cytometry analysis of number of dead cells of the indicated species as shown in A. Colonies were grown on TSB medium supplemented with glucose (1% W/V), and TSB medium supplemented with glucose (1% W/V) + buffer. Data were collected from 72 hours post inoculation, 100,000 cells were counted. Y-axis represents the % of dead cells, graphs represent mean ± SD from three independent experiments (n = 9). Statistical analysis was performed between glucose and glucose + buffer using unpaired two tailed t test with Welch’s correction. P < 0.05 was considered statistically significant. **D)** Table showing multiple comparison test between dead cell populations of indicated species grown in TSB medium supplemented with glucose (1% W/V). Statistical analysis was performed using Brown-Forsthye and Welch’s ANOVA with Dunnett’s T3 multiple comparisons test. P < 0.05 was considered statistically significant.

To better assess the level of stress on the single-cell level we grew the bacteria on solid growth media (TSB), TSB with glucose, TSB with raffinose in the presence and absence of a buffering system. Colony cells grown without glucose are rod-shaped, divide normally, and are arranged in short chains (Figure 5). While all the species looked similar in TSB, glucose-induced noticeable alterations in the morphology of *LGG* and *L. paracasei* cells. Noticeable clumps were induced in both species, which also lost their characteristic elongated rod shape. Alteration in cell shape upon glucose treatment could be partially rescued with the application of buffer, supporting a differential response of probiotic bacteria to fermentation and its acidic products. Interestingly, alterations in cell shape did not perfectly correlate with the response to acid stress as judged by cell growth as LGG had pronounceable growth upon utilization of glucose, while *L. paracasei* failed to increase it carrying capacity with glucose. Therefore, the alterations in cell shape may be independent adaption to carbohydrate (for example a change in surface proteome) and not a direct reflection of acid stress.

**Figure 5.**
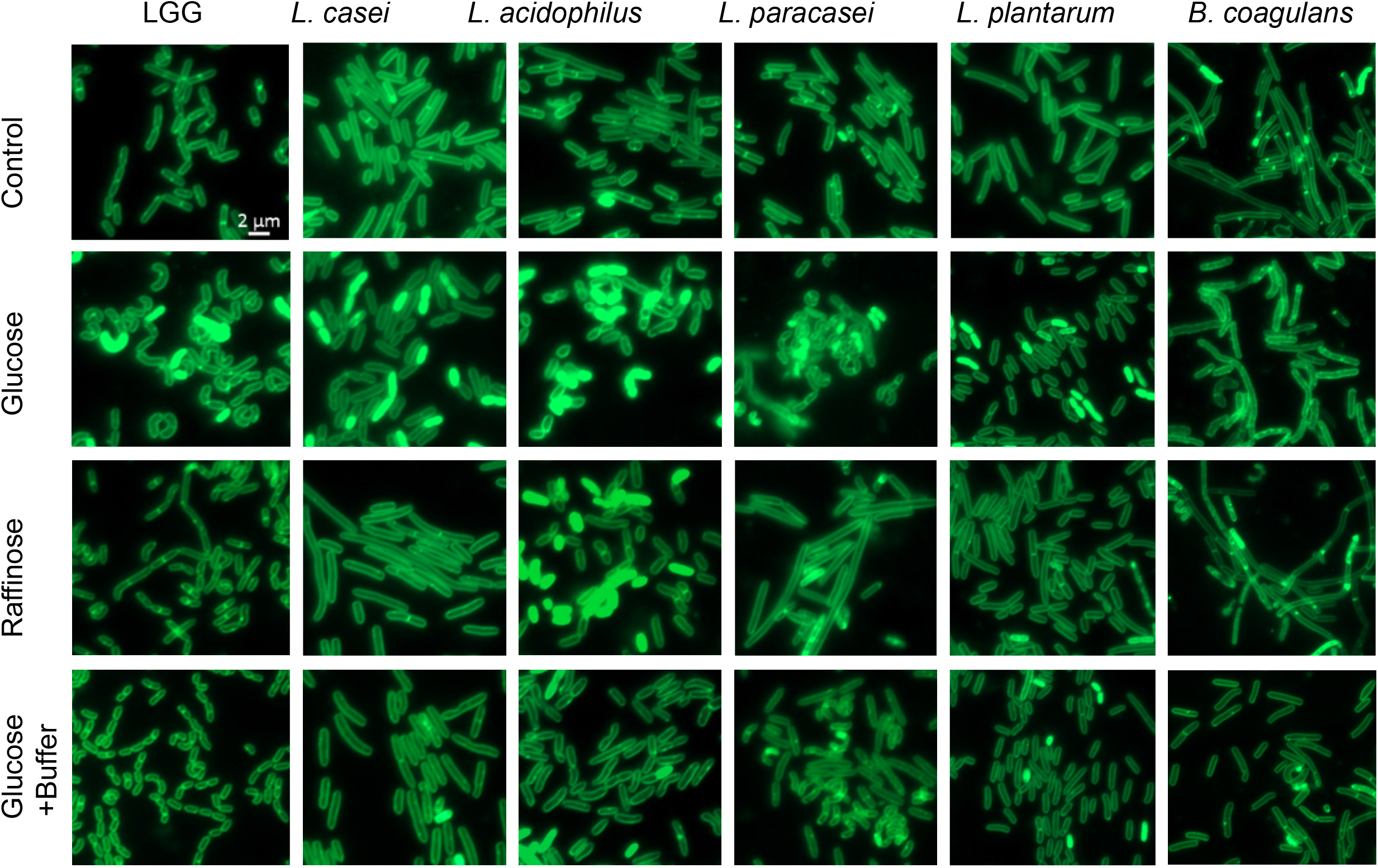
Self-imposed acidic stress induces a polymorphism in cellular arrangements of colony cells. Shown are the indicated fluorescence microscope images of cells from biofilm colonies that were grown on TSB medium (control), TSB medium supplemented with glucose and raffinose (1% W/V), and TSB medium supplemented with glucose (1% W/V) + buffer. Cells were stained using Green-membrane stain FM™ 1-43FX. Biofilms were grown at 37° C in CO_2_ enriched environment. Cells were imaged at 72 hours post inoculation. The images represent 3 independent experiment, from each repeat at least 10 fields examined. Scale bar = 2µm.

## Discussion

Probiotic bacteria are considered a means for microbiota modulation, along with nutrition, personal hygiene, and lifestyle (1). Probiotics are found to have a beneficial effect on the consumer in the prevention of antibiotic-associated diarrhea and acute infectious diarrhea, treatment of inflammatory bowel disease (26), and other gastric disorders(27). Lactobacillaceae are widely used probiotic bacteria, represented in most fermented products and supplements. Probiotic performance in the gut is dependent on nutrient composition and availability (28). Positive effects on probiotics proliferation and beneficial effects on the host have been associated with prebiotic consumption. For example, it was shown that the addition of raffinose and *L. acidophilus* to the diet of rats decreases their body weight and increases the concentration of *L. acidophilus* in the gut (29). However, the responses of single LAB strains to their own acidic fermentation products may vary and contribute to their physiological responses to carbohydrates.

Therefore, we systematically compared five different LAB species for their response to utilization of glucose and raffinose model sugars. We found that glucose utilization has a general role in shaping the colony morphology under static conditions, as all species exhibit morphological colony changes upon glucose treatment. Similarly, medium acidification was generally similar during static growth (Figure 4B) and comparable for all Lactobacillaceae during planktonic growth (Figure 2). For all tested species, the final product of glucose utilization and fermentation (occurring under microaerophilic conditions) was almost exclusively the organic acid lactate alongside acetate. Analyses of fermentation capacities of the different species revealed similar fermentation efficiencies. Lactic and acetic acid concentrations and the pH of the growth medium were all similar and stable among the different species (Figures 3 and 4B). In agreement, the alignment of lactate dehydrogenase proteins revealed that *LGG, L. paracasei* and *L. casei* have two groups of almost identical protein homologs with more than ninety percent identity between the different species (Figure S3).

In contrary to glucose utilization, organic acid accumulation, and medium acidification that are comparable between strains, the carrying capacity of the culture is differentially enhanced during planktonic growth of Lactobacillaceae. While *LGG, L. acidophilus L. casei* and *L. plantarum* doubled the carrying capacity following exposure to glucose exposure, *L. paracasei* had no noticeable growth induction with glucose. Interestingly, the capacity of LAB strains to utilize glucose, raffinose, mannose, and xylose for planktonic growth differed significantly (Figures 1, 2, and S1). A detailed analysis of the microbial growth reflected that carbon utilization is unintuitively inert to the growth rate, and does not affect the lag phase (Figure 2). Rather, it allows the bacterial community to reach a significantly higher carrying capacity that may account for changes in their proportions in the GI.

Acid stress in Lactobacillaceae is self-imposed stress, and, thus, LAB are relatively acid-tolerant, and employ several mechanisms to regulate the homeostasis of the pH level (30). A. The mechanisms generally include the removal of protons or alkalization of the environment via ammonia production through the arginine deiminase (ADI) (31). In addition, glutamic acid decarboxylase (GAD) catalyzes the decarboxylation of glutamate into gamma-aminobutyric acid (GABA), which results in the alkalization of the cytoplasmic pH due to the removal of protons (32). Lastly, the urease system allows the hydrolysis of urea, which enhances the survival of LAB under acid stress conditions by the production of NH_3_. The urease operon was found to be positively regulated under low pH levels in the LAB *Streptococcus salivarius* (33). F-ATPase system is another mechanism that protects LAB from acid damage. Overall, probiotic acid-tolerant species are of great value as additional encapsulation in order to ensure survival to transit through the acidic stomach. As the survival of LAB during acid stress is quite heterogeneous the expression of these systems during self-imposed acid stress and, in the stomach, needs to be properly evaluated while selecting formulating probiotics. Alternatively, food carriers with high buffering potential to aid gastric transit and ensure that viable cells reach the small intestine may contribute to assure a beneficial effect on the host. For example, our results indicate that *L. paracasei* in liquid (Figure 1) and *L. acidophilus* (Figure 4) grown on a solid medium exhibit poor acid tolerance during self-imposed acid stress. One immediate application of our findings for *L. paracasei* in solution (Figure 1) and *L. acidophilus* (Figure 4C) is that probiotic performance of these strains in the gut is dependent in the availability of a suitable buffering system.

In the gut, acid-sensitive species are protected from the acidic pH of the stomach (2-4) because of the buffering properties of food, which depends on the type of food and its volume (34, 35). In parallel, self-imposed acid stress from accumulation of organic acids following the utilization of carbohydrates may affect the performance of the ingested bacterial species. Interestingly, buffering of the colonies restored glucose-free morphology of all strains, indicating that acid stress is involved in biofilm formation throughout the Firmicutes phylum directly or indirectly. The addition of a buffering system to the growth media of bacteria grown in the presence of glucose increased the survival of most species significantly. However, while cell death was decreased with buffering, it was still significantly higher with glucose compared with non-glucose conditions (Figure 4C). Thus, the glucose-induced change in colony morphology may be reflective of a complex adaptation and biofilm regulation, rather than a simple reflection of cell density. While glucose utilization capacities and medium acidification under our conditions were similar, the enhancement of carrying capacity (Figure 2), carbohydrate mediated cell death within the colony (Figure 4C), and the adaptation to glucose utilization at the single-cell level (Figure 5) varied dramatically between the tested species. Changes in the surface proteome(14), quorum sensing(36), and the activation of differential target genes of the stressosome (37) may differ in a context-dependent manner between LAB strains. While the exact regulations remain to be determined, our results indicate that the overall physiological response to carbohydrate metabolism and self-imposed acid stress is not a mare reflection of microbial growth.

## Materials and methods

### Strains, Media and imaging

*Lactobacillus acidophilus* ATCC 4356, *Lacticaseibacillus casei* ATCC 393, *Lacticaseibacillus casei subsp. paracasei* ATCC BAA-52, *Lactiplantibacillus plantarum* ATCC 8014 and *Lacticaseibacillus rhamnosus GG* ATCC probiotic strains, were used in the study. *Bacillus coagulans* ATCC 10545 was used as a control. A single colony of Lactobacillaceae were isolated on a solid deMan, Rogosa, Sharpe Agar (MRS) plate, was inoculated into 5ml MRS broth (Difco, Le Pont de Claix, France), and grown at 37°C, without shaking, for overnight. A single colony of *Bacillus coagulans* isolated on a solid LB agar plate was inoculated into 5ml LB broth (Difco) and grown at 37°C, with shaking overnight. For biofilm colonies, these cultures were inoculated into a solid medium (1.5% agar) contains 50% Tryptic soy broth (TSB), TSB supplemented with (1% w/v) D-(+) - glucose, (1% w/v) D-(+)- raffinose or (1% w/v) D-(+)- mannose or (1% w/v) D-(+)-Xylose or with (1% w/v) D-(+) - glucose buffered with MOPS and potassium phosphate buffer. The bacteria were incubated in a BD GasPak™ EZ - Incubation Container with BD GasPak EZ CO2 Container System Sachets (260679) (Becton, Sparks, MD, USA), for 72 hours in 37°C. All images were taken using a Stereo Discovery V20” microscope (Tochigi, Japan) with objectives Plan Apo S ×1.0 FWD 60 mm (Zeiss, Goettingen, Germany) attached to a high-resolution microscopy Axiocam camera. Data were created and processed using Axiovision suite software (Zeiss). For planktonic growth, the bacterial cultures were inoculated into a liquid medium 50% Tryptic soy broth (TSB) with different sugars as described above, incubated for 24h, at 37°C.

### Growth measurement and analysis

Cultured cells grown for overnight were diluted 1:100 in 200 μl liquid medium contains 50% Tryptic soy broth (TSB) (BD), TSB supplemented with (1% w/v) D-(+)- glucose, (1% w/v) D-(+)- raffinose or (1% w/v) D-(+)- mannose or or (1% w/v) D-(+)-Xylose in a 96-well microplate (Thermo Scientific, Roskilde, Denmark). Cells were grown with agitation at 37 °C for 18 h in a microplate reader (Tecan (Infinite® 200 PRO, Switzerland), and the optical density at 600 nm (OD_600_) was measured every 30 min. Maximum OD, growth rate and lag time calculation were performed with GrowthRates 3.0 software.

### Fluorescence microscopy

A bacterial colony grown as described above was suspended in 200 μl 1x Phosphate-Buffered Saline (PBS), and dispersed by pipetting. Samples were centrifuged briefly, pelleted and re-suspended in 5μl of 1x PBS supplemented with the membrane stain FM1-43 (Molecular Probes, Eugene, OR, USA) at 1 μg/ml. This cells were placed on a microscope slide and covered with a poly-L-Lysine (Sigma) treated coverslip. The cells were observed by Axio microscope (Zeiss, Germany). Images was analyzed by Zen-10 software (Zeiss).

### pH measurements

After inoculation in different liquid mediums and 24h incubation (without shaking) at 37°C, the cells were separated from the medium by centrifugation (4000xg, 20 min) followed by filtration through a 0.22 μm. The conditioned media acidity was measured by the pH meter (Mettler Toledo). pH measurements of the solid media were done using pH-indicator strips (MQuant®, Merck).

### Determination of organic acids

The supernatant of the bacteria grown on MSgg with 1% glucose for 24h was filtered through 0.22 μm filter membranes for HPLC analysis. The contents of organic acids in each liquid sample were determined using an Infinity 1260 (Agilant Technologies) HPLC system with C 18 column (Syncronis™ C18 Columns, 4.6 × 250 mm, 5 μm). Mobile phase A was acetonitrile and mobile phase B was 5-mM KH_2_PO_4_ pH 2.4. The flow rate was kept constant at 1 mL/min, with ultraviolet detection performed at 210 nm. The injection volume was 20 μL and the column temperature was maintained at 30 °C. Identification and quantification of organic acids were accomplished by comparing the retention times and areas with those of pure standards. Notably, although quantification of organic acids in TSB was not feasible due to high background, the comparable acidity strongly indicates similar fermentation capacities of the strains tested in rich growth media.

### Flow Cytometry Analysis

Starter cultures were spotted on TSB with glucose or TSB with glucose buffered with MOPS and potassium phosphate buffer. The plates were then incubated as mentioned in the first section. Colonies were harvested after 72 hours. Samples were diluted in PBS and measured using an LSR-II new cytometer (Becton Dickinson, San Jose, CA, USA). PI (Propidium Iodide, 20Mm, Invitrogen™) fluorescence was measured using laser excitation of 488 nm, coupled with 600 LP and 610/20 sequential filters. A total of 100,000 cells were counted for each sample and flow cytometry analyses was performed using BD FACSDiva™ software.

### Protein alignment, phylogenetic tree

Phylogenetic tree was built based on 16S of the bacteria and multiple alignment sequences of LDH proteins was preformed using Clustal Omega. https://www.ebi.ac.uk/Tools/msa/clustalo/

### Statistical analysis

All experiments were performed at least three separate and independent times in triplicates, unless stated otherwise. Statistical analyses were performed with GraphPad Prism 9.0 (GraphPad 234 Software, Inc., San Diego, CA). Relevant statistical test are mentioned in the indicated legends of the figures.

## Supporting Information

**Figure S1.**
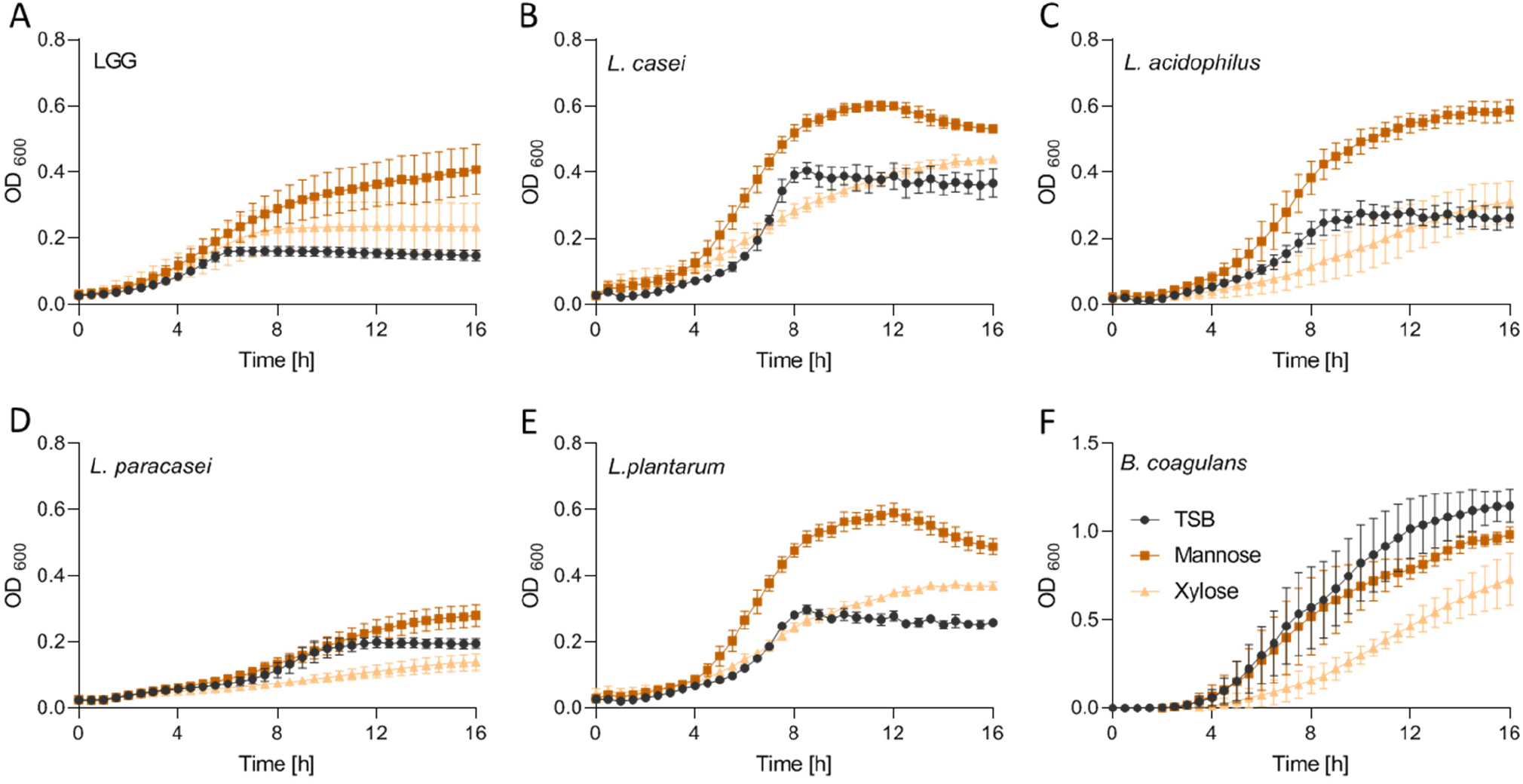
The effect of fermentable sugars on planktonic growth. Planktonic growth of the indicated species (A) LGG (B) *L. casei* (C) *L. acidophilus* (D) *L. paracasei* (E) *L. plantarum* (F) and *B. coagulans* in TSB medium (control) and TSB medium supplemented with mannose (1% W/V) and xylose (1% W/V). Graphs represent mean ± SD from three independent experiments (n =3).

**Figure S2.**
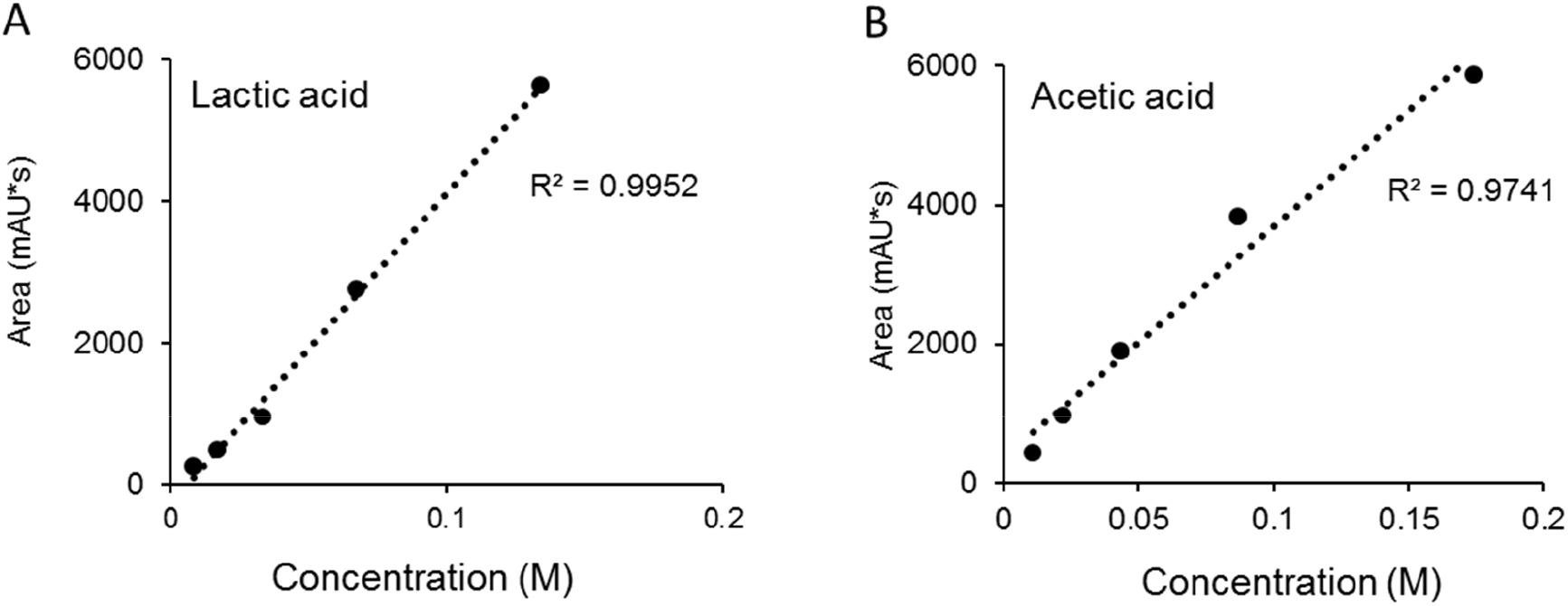
**Calibration curves** for lactic acid (A) and acetic acid (B)

**Figure S3.**
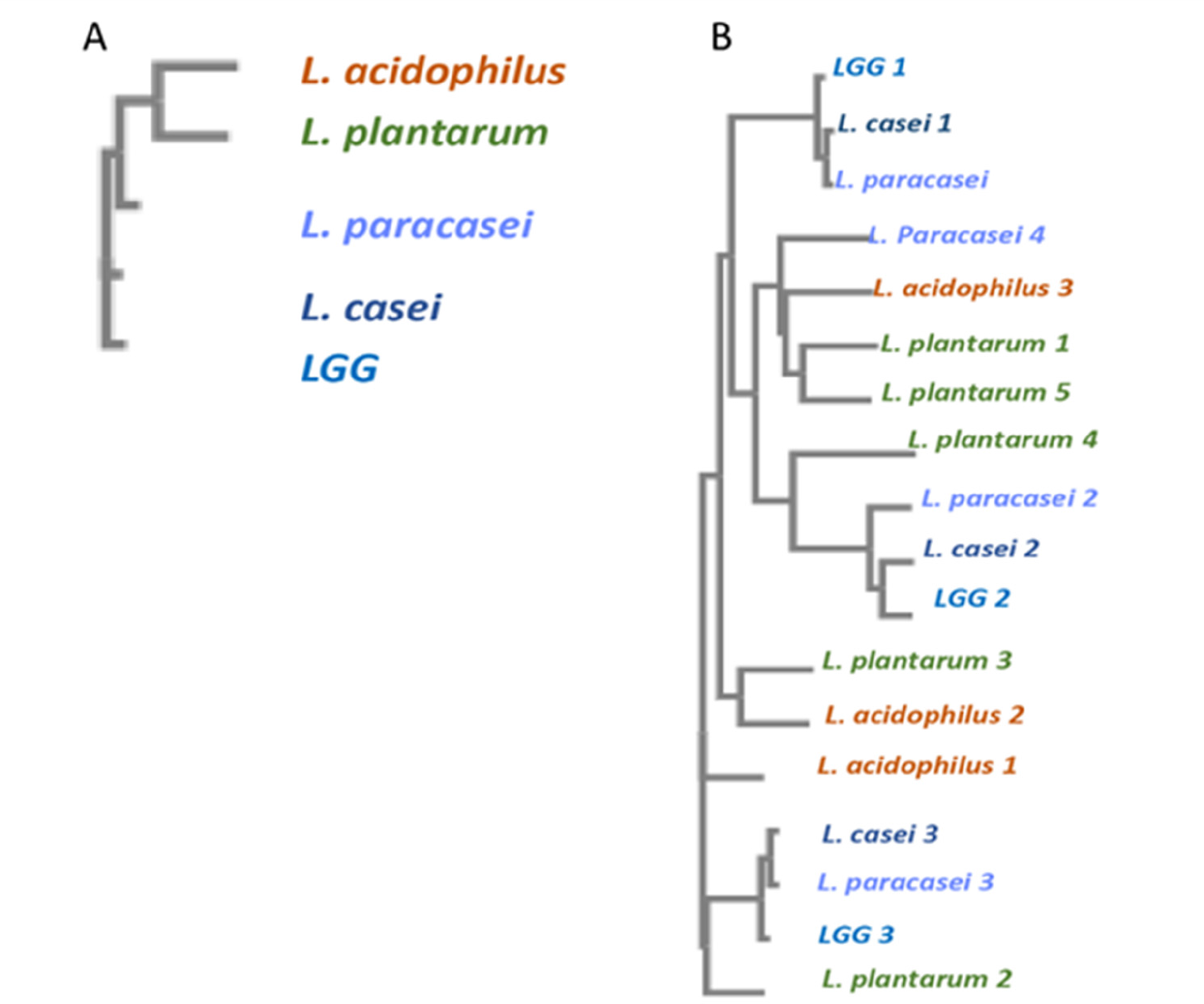
In silico analysis of LDH proteins. Phylogenetic trees were generated using Clustal Omega based on A)16S or B) L- LDH proteins

**Table S1.** Table shows the individual values of pH measurements indicated in figure 4B from three independent experiments.

|  | TSB | TSB | TSB | Glucose | Glucose | Glucose | Raffinose | Raffinose | Raffinose |
| --- | --- | --- | --- | --- | --- | --- | --- | --- | --- |
| LGG | 6 | 6 | 6 | 4 | 4 | 4 | 6 | 6 | 6 |
| *L. casei* | 6 | 6 | 6 | 4 | 4 | 4 | 5 | 5 | 5 |
| *L. acidophilus* | 6 | 6 | 6 | 4 | 4 | 4 | 5 | 5 | 5 |
| *L. paracasei* | 6 | 6 | 6 | 4 | 4 | 4 | 5 | 5 | 5 |
| *L. plantarum* | 6 | 6 | 6 | 4 | 4 | 4 | 5 | 5 | 5 |
| *B.coagulans* | 6 | 6 | 6 | 4.5 | 4.5 | 5 | 5.5 | 5.5 | 5.5 |

